# Neuroinflammation and protein aggregation co-localize across the frontotemporal dementia spectrum

**DOI:** 10.1101/525642

**Authors:** W Richard Bevan-Jones, Thomas E Cope, P Simon Jones, Luca Passamonti, Young T Hong, Tim D Fryer, Robert Arnold, Jonathan P Coles, Franklin I Aigbirhio, Andrew J Larner, Karalyn Patterson, John T O’Brien, James B Rowe

## Abstract

The clinical syndromes of frontotemporal dementia are clinically and neuropathologically heterogeneous, but processes such as neuroinflammation may be common across the disease spectrum. We investigated how neuroinflammation relates to the aggregation of Tau and TDP-43 in frontotemporal dementia, and to the heterogeneity of clinical disease. We used positron emission tomography *in vivo* with (a) [^11^C]PK-11195, a marker of activated microglia and a proxy index of neuroinflammation, and (b) [^18^F]AV-1451, a radioligand with increased binding to pathologically affected regions in tauopathies and diseases associated with TDP-43 protein aggregation, and which is used as a surrogate marker of non-β-amyloid protein aggregation. We assessed 31 patients with frontotemporal dementia (10 with behavioural variant frontotemporal dementia, 11 with the semantic variant of primary progressive aphasia and 10 with the non-fluent variant of primary progressive aphasia), 28 of whom underwent both [^18^F]AV-1451 and [^11^C]PK-11195 PET, and matched controls (14 for [^18^F]AV-1451 and 15 for [^11^C]PK-11195). We used univariate region-of-interest analyses, and multivariate analysis of the distribution of binding that explicitly control for individual differences in ligand affinity for TDP-43 and different Tau isoforms. We found differences between patients and controls in frontotemporal regions for both neuroinflammation and protein aggregation, and a strong positive correlation between these two processes in all disease groups. Despite this regional co-localisation, the multivariate distribution of [^11^C]PK-11195 binding related better to clinical heterogeneity than did the distribution of [^18^F]AV-1451: distinct spatial modes of neuroinflammation were associated with different frontotemporal dementia syndromes and supported accurate group classification of participants. These *in vivo* findings indicate a close association between neuroinflammation and protein aggregation in frontotemporal dementia. The inflammatory component may be important in shaping the clinical and neuropathological patterns of the diverse clinical syndromes of frontotemporal dementia.

## Introduction

Frontotemporal dementia (FTD) encompasses a clinically and pathologically heterogeneous group of neurodegenerative conditions, including the behavioural variant (bvFTD) (Rascovsky et al., 2011), non-fluent variant primary progressive aphasia (nfvPPA) and semantic variant primary progressive aphasia (svPPA) (Gorno-Tempini et al., 2011). In recent years, attention has focused on understanding the pathogenic role of protein misfolding and aggregation, which is a cardinal feature of the *post mortem* diagnostic criteria for frontotemporal lobar degeneration (MacKenzie et al., 2010). However, there are several different pathological proteins and aggregation morphologies in FTD, with generally weak correlations between clinical syndrome and the type of pathological protein (Seelaar, Rohrer, Pijnenburg, Fox, & van Swieten, 2011) (with the exception of svPPA, which is strongly associated with TDP-43 type C neuropathology (Spinelli et al., 2017)). However, other neuropathological processes may be present in common across these diverse clinical syndromes and present potential therapeutic targets. In particular, there is converging evidence for the role for neuroinflammation in neurodegenerative dementias, including FTD, from genetic associations (Broce et al., 2018; Guerreiro et al., 2013; Rayaprolu et al., 2013), cerebrospinal fluid (Sjogren, Folkesson, Blennow, & Tarkowski, 2004; Woollacott et al., 2018), epidemiology (Miller et al., 2013, 2016), *post mortem* tissue (Lant et al., 2014; Venneti, Wang, Nguyen, & Wiley, 2008) and animal models (Bhaskar et al., 2010; Yin et al., 2010; Yoshiyama et al., 2007). Both the intensity of neuroinflammation, and its distribution across the brain, may be relevant determinants of the clinical syndrome.

Positron emission tomography (PET) allows the topographic quantification of specific molecules using radioligands. In this study, we measured neuroinflammation and protein aggregation *in vivo* in patients with bvFTD, svPPA and nfvPPA, to answer key questions regarding the relationship of these pathophysiological processes. [^11^C]PK-11195, which binds to the translocator protein (TSPO) that is expressed on the outer mitochondrial membrane of activated microglia, is a robust and sensitive marker of microglial activation with an established role as a proxy for neuroinflammation in neurodegenerative diseases (Stefaniak & O’Brien, 2015). [^18^F]AV-1451 was originally developed to bind to paired helical filament Tau in Alzheimer’s disease (Chien, Bahri, Szardenings, Walsh, & Mu, 2013; Xia et al., 2013; W. Zhang et al., 2012), and has been extensively used in Alzheimer’s and non-Alzheimer’s diseases. Elevated *in vivo* binding is seen in tauopathies characterised by straight filaments (W. Richard Bevan-Jones et al., 2016; Jones et al., 2018; Luca Passamonti et al., 2017; Smith et al., 2017), albeit with generally lower binding affinity than in Alzheimer’s disease, and also in TDP-43 related disease (R. W. Bevan-Jones et al., 2018; William Richard Bevan-Jones et al., 2017; Makaretz et al., 2017). It also has low affinity for beta-amyloid and alpha-synuclein (Xia et al., 2013). Therefore, although the molecular interpretation of increased binding is incompletely understood (Marquié et al., 2015; Sander et al., 2016), this elevated *in vivo* binding suggests [^18^F]AV-1451 may represent a proxy index of aggregated non-β-amyloid pathological proteins across the FTD spectrum.

Given the evidence for differences in affinity of [^18^F]AV-1451 for different Tau and TDP-43 conformational targets, our analysis strategy concentrates on the relative topographical distribution of binding across regions within each individual, rather than the simple magnitude of binding. In this way we explicitly control for difference in binding affinity between syndromes and protein strains within each syndrome.

We test the hypotheses that, in FTD, neuroinflammation and protein aggregation are both increased in frontotemporal regions compared to controls, and that neuroinflammation and protein aggregation co-localise in each FTD syndrome, consistent with the syndrome-specific neuropathological distributions (e.g., co-localization of neuroinflammation and protein aggregation in the temporal pole of patients with svPPA). We use data driven approaches to elucidate the spatial modes of neuroinflammation associated with FTD, and machine learning based on multi-dimensional scaling of distributional dissimilarities, to investigate whether the cortical distribution of neuroinflammation and protein aggregation can accurately discriminate diagnostic groups thereby illustrating their mechanistic importance.

## Materials and methods

As part of the NIMROD study (W Richard Bevan-Jones et al., 2017), 31 patients (10 with bvFTD, 11 with svPPA and 10 with nfvPPA) underwent PET scanning with [^18^F]-AV1451. 28 of the 31 (9 with bvFTD, 9 with svPPA and 10 with nfvPPA) also underwent a PET scan with [^11^C]PK-11195. The order of scans was randomised. Fourteen healthy control participants underwent [^18^F]AV-1451 PET and, to minimise radiation exposure in healthy individuals, a different group of 15 healthy participants underwent [^11^C]PK-11195 PET scanning. Genetic and amyloid status (by PET or cerebrospinal fluid biomarkers) for patients were tested if clinically indicated.

PET with [^18^F]AV-1451 and [^11^C]PK-11195 was performed on a GE Discovery 690 PET/CT (GE Healthcare) with a low dose CT for attenuation correction or on a GE Advance PET scanner (GE Healthcare) with a 15-min 68Ge/68Ga transmission scan for attenuation correction. The PET scan itself used dynamic imaging for 90 ([^18^F]AV-1451) and 75 ([^11^C]PK-11195) minutes respectively. All radioligands were prepared at the Wolfson Brain Imaging Centre (WBIC), University of Cambridge, with high radiochemical purity (>95%). Each subject underwent contemporaneous 3T magnetic resonance imaging (MRI) using a Siemens Magnetom Skyra, Verio or Tim Trio (www.medical.siemens.com). A high-resolution T1 weighted sequence was acquired (176 slices of 1.0 mm thickness, TE= 2.98 ms, TR = 2300 ms, flip angle =9°, acquisition matrix 256×240; voxel size = 1×1×1 mm3) and used for tissue segmentation (grey and white matter along with CSF), and for non-rigid registration of standard space regions of interest. For both ligands, non-displaceable binding potential (BP_ND_) was calculated in 83 regions of interest, defined by a Hammers atlas modified to include the midbrain and the dentate nucleus of the cerebellum, by kinetic modelling using a simplified reference tissue model, with cerebellar grey matter as reference region for [^18^F]AV-1451 (Luca Passamonti et al., 2017) and supervised cluster analysis used to define the [^11^C]PK-11195 reference region (Yaqub et al., 2012). Prior to kinetic modelling all region of interest data were corrected for cerebrospinal fluid contamination of the region (i.e. partial volume corrected) through division by the mean region grey plus white matter fraction, determined using tissue probability maps smoothed to PET spatial resolution.

Four data analysis approaches were used, each designed to answer a different focused question and to explicitly control for expected between-subject and between-region differences in ligand affinity.

As a first-stage data exploration of between-group differences, a repeated-measures ANOVA was performed across the 83 regions, including age as a covariate and Greenhouse-Geisser penalisation of degrees of freedom to correct for non-sphericity. *Post hoc* t-tests were then performed between each group, corrected for false discovery rate (FDR) over regions.

Second, to examine the relationship between neuroinflammation and protein aggregation in each disease group, a correlation between the regional BP_ND_ of each ligand was performed. PET scanning with any ligand characteristically results in a general pattern of lower BP_ND_ in brain regions such as temporal lobe and higher BP_ND_ in deep brain nuclei. We were concerned that such non-specific effects might drive apparent correlations, and weak correlations were observed between our cohorts of controls for each ligand (supplementary figure 1). To control for this, we examined the between-ligand correlation within each disease group both with and without subtraction of the control mean BP_ND_ for each of the 83 regions of interest.

**Figure 1:**
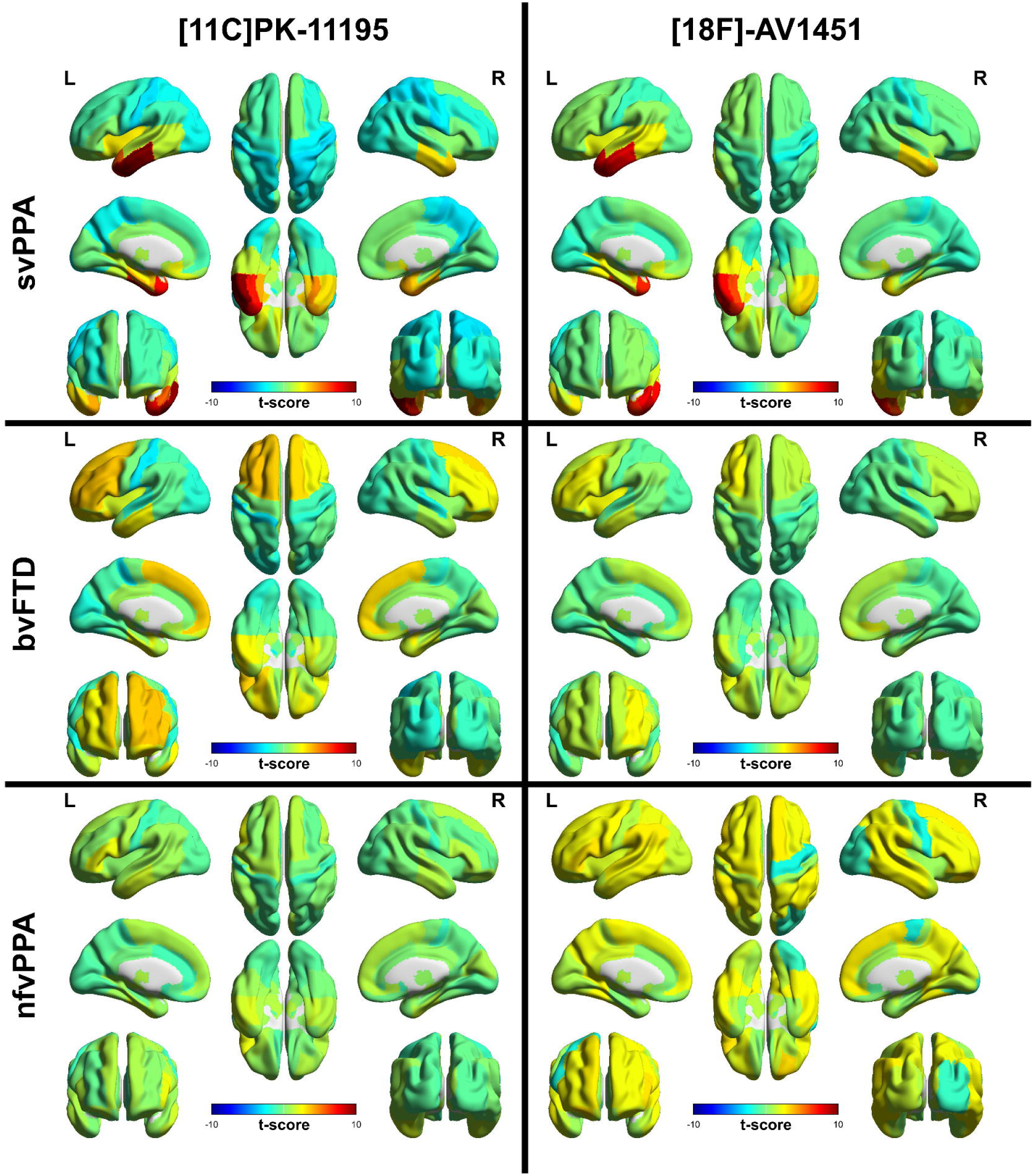
Unthresholded regional T-scores for each disease group compared to the control group for [^11^C]PK-11195 BP_nd_ in the left column and [^18^F]AV-1451 BP_ND_ in the right column.

Third, to elucidate the topographical patterns of inflammation and protein aggregation in FTD, we entered the BP_ND_ of each ligand in each of the 83 regions of interest into a principal component analysis. Components were retained by Cattell’s criterion (i.e. to the elbow of the Scree plot) and then tested for group differences across diagnosis in a repeated measures ANOVA. *Post hoc* t-tests examined group differences in the expression of each topographical pattern. These first three analyses were performed in SPSS Statistics version 25 (IBM).

Finally, we undertook an analysis of the relative distribution of ligand binding potential for each ligand for every individual. This used previously published non-parametric methods (W. Richard Bevan-Jones et al., 2016), that were explicitly designed to control for between-subject differences in the scaling of each ligand, such as might result from differences in the affinity of [^18^F]AV-1451 for different conformations of Tau or TDP-43, as well as spatial dependence between adjacent regions in PET data due to signal spread. These methods can be conceptualised as analogous to multi-voxel pattern analysis techniques for fMRI (Kriegeskorte, Mur, & Bandettini, 2008), but rather than attempting to classify observed stimuli within an individual on the basis of their representational similarity, here we are attempting to classify individuals on the basis of the similarity of relative ligand BP_ND_ distributions within their brain, blinded to overall differences in binding affinity. To do this, for each ligand and every individual separately, the parcellated data were converted to 83-element linear vectors. For each ligand separately, the resultant vectors were non-parametrically correlated (Spearman’s rho) pairwise between individuals, resulting in two matrices that represented the similarity of each individual’s scan to each other individual for that ligand. The inverse of these matrices (i.e. the between-individual dissimilarities) were used to calculate a two-dimensional scaling for each disease sub-group pair, using the squared metric stress distance criterion of the ‘mdscale’ function in Matlab R2017b (Mathworks). The resulting locations in two-dimensional space formed the inputs to a ten-fold crossvalidated linear support vector machine (CV-SVM) for between-group classification based on each ligand separately. Statistical significance of the classification was assessed by comparison of the loss function of the CV-SVM against a null distribution of loss functions created by 1000 repetitions of the same procedure for identical data but shuffled group assignment labels. For those individuals who underwent scanning with both ligands, the CV-SVM process was repeated on multi-modal, four-dimensional scaling.

## Results

Summary demographics are outlined in table 1, and neuropsychological test scores, motor features, genetic and CSF status for each participant are provided in table 2. Within the bvFTD group, two patients were positive for pathogenic mutations in the microtubule associated protein tau (MAPT) and three for expansions in C9 open reading frame 72 (C9orf72). One of the nfvPPA group had a mutation in progranulin (GRN). CSF or PET amyloid status was assessed in 6 participants (4 with svPPA, and 2 with nfvPPA), all of whom were negative.

**Table 1:**
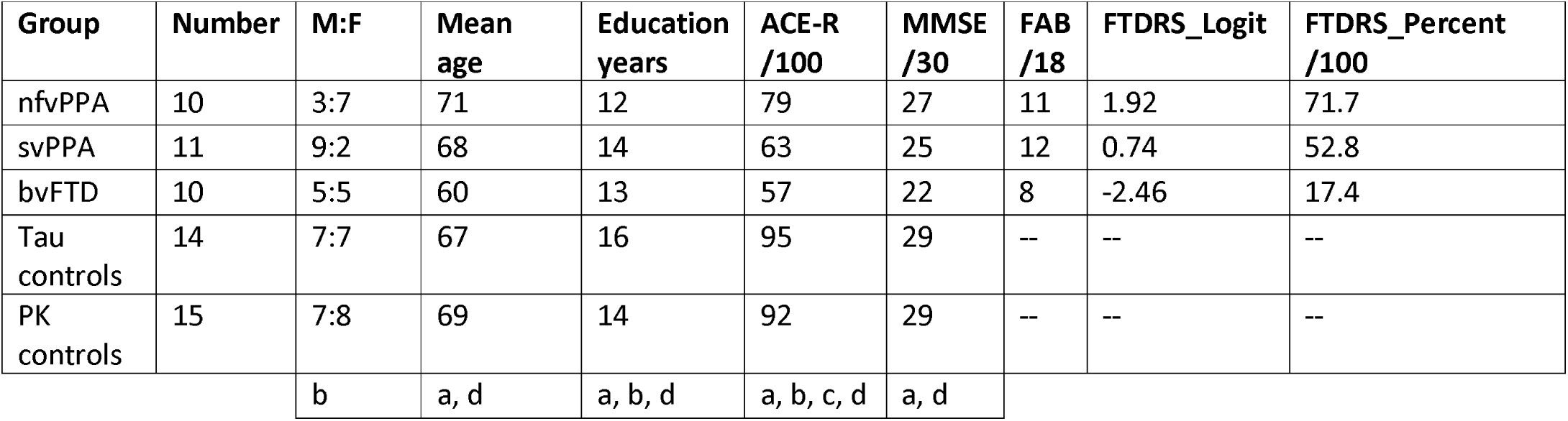
Summary demographics and neuropsychometry: a=F test significant p<0.05 across all groups, b=p<0.05 significant pairwise comparison nfvPPA vs combined control group, c=p<0.05 significant pairwise comparison svPPA vs combined control group, d=p<0.05 significant pairwise comparison bvFTD vs combined control group. Pairwise comparisons are by t-test for each demographic except sex comparison by Chi-squared.

| Group | Number | M:F | Mean age | Education years | ACE-R /100 | MMSE /30 | FAB /18 | FTDRS_Logit | FTDRS_Percent /100 |
| --- | --- | --- | --- | --- | --- | --- | --- | --- | --- |
| nfvPPA | 10 | 3:7 | 71 | 12 | 79 | 27 | 11 | 1.92 | 71.7 |
| svPPA | 11 | 9:2 | 68 | 14 | 63 | 25 | 12 | 0.74 | 52.8 |
| bvFTD | 10 | 5:5 | 60 | 13 | 57 | 22 | 8 | -2.46 | 17.4 |
| Tau controls | 14 | 7:7 | 67 | 16 | 95 | 29 | -- | -- | -- |
| PK controls | 15 | 7:8 | 69 | 14 | 92 | 29 | -- | -- | -- |
|  |  | b | a, d | a, b, d | a, b, c, d | a, d |  |  |  |

**Table 2:**
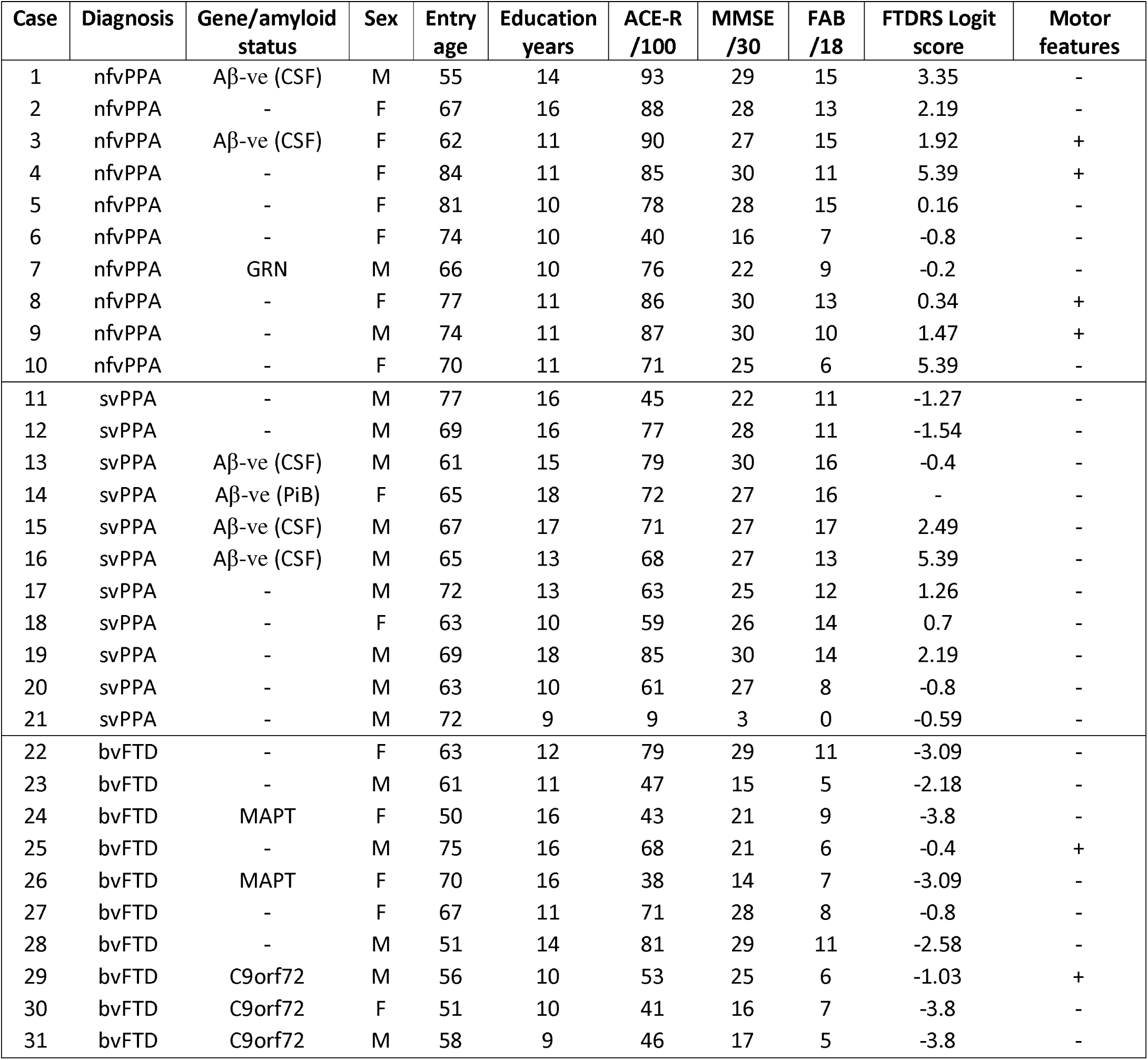
Demographics, neuropsychological testing, genetic/amyloid status and motor phenotype for each disease participant. Aβ-ve: negative tests for beta-amyloid by cerebrospinal fluid biomarkers (CSF) or Pittsburgh compound B PET scan. MAPT: microtubule associated protein tau.

### Group comparisons of frontotemporal dementia with controls

The repeated-measures ANOVA of regional [^11^C]PK-11195 binding across the FTD groups and controls demonstrated a significant interaction between region and diagnosis (F(39.5, 500.6)=9.2, p<0.0001). T-maps from the post-hoc pairwise comparisons between the control group and each disease group are shown in figure 1. After correction for FDR, regions with significantly elevated binding were: in the bvFTD group; bilateral superior frontal gyri and putamen, right nucleus accumbens, left posterior orbital gyrus, inferior frontal gyrus and middle frontal gyrus. In the svPPA group; left insula, middle and inferior temporal gyri, right superior parietal gyrus, middle and inferior temporal gyrus, bilateral postcentral gyri, superior temporal gyrus, parahippocampal and ambient gyri, amygdala, inferior lateral anterior temporal lobe, medulla, nuclei accumbens, medial anterior temporal lobe, fusiform gyri. In the nfvPPA group no differences survived FDR correction but the peak t-score was in left inferior frontal gyrus (t(23)=2.17, uncorrected p=0.04), which would be expected *a priori* to be the disease epicentre (Rogalski et al., 2011).

The repeated measures ANOVA of regional [^18^F]AV-1451 binding across the FTD groups and controls showed a significant interaction between region and diagnosis (F(33.4, 445.1)=10, p<0.0001). T-maps from the post-hoc pairwise comparisons between the control group and each disease group are shown in figure 1. After correction for FDR, significantly elevated binding was seen in svPPA, in the following regions; left amygdala, fusiform, medial anterior temporal lobe, middle and inferior temporal gyri and superior temporal gyrus, bilateral inferolateral anterior temporal lobes.

### Correlation of [^11^C]PK-11195 with [^18^F]AV-1451 in frontotemporal dementia

Regional control adjusted group mean [^11^C]-PK11195 BP_ND_ was strongly correlated with regional group mean [^18^F]-AV1451 BP_ND_ in each group both before and after the subtraction of the control group values in every region: svPPA (r(81) = 0.727, p<0.0001 before, r(81) = 0.883, p<0.0001 after), bvFTD (r(81) = 0.582, p<0.0001 before, r(81) = 0.499, p<0.0001 after), and nfvPPA (r(81) = 0.427, p<0.0001 before, r(81) = 0.589, p<0.0001 after) (figure 2).

**Figure 2:**
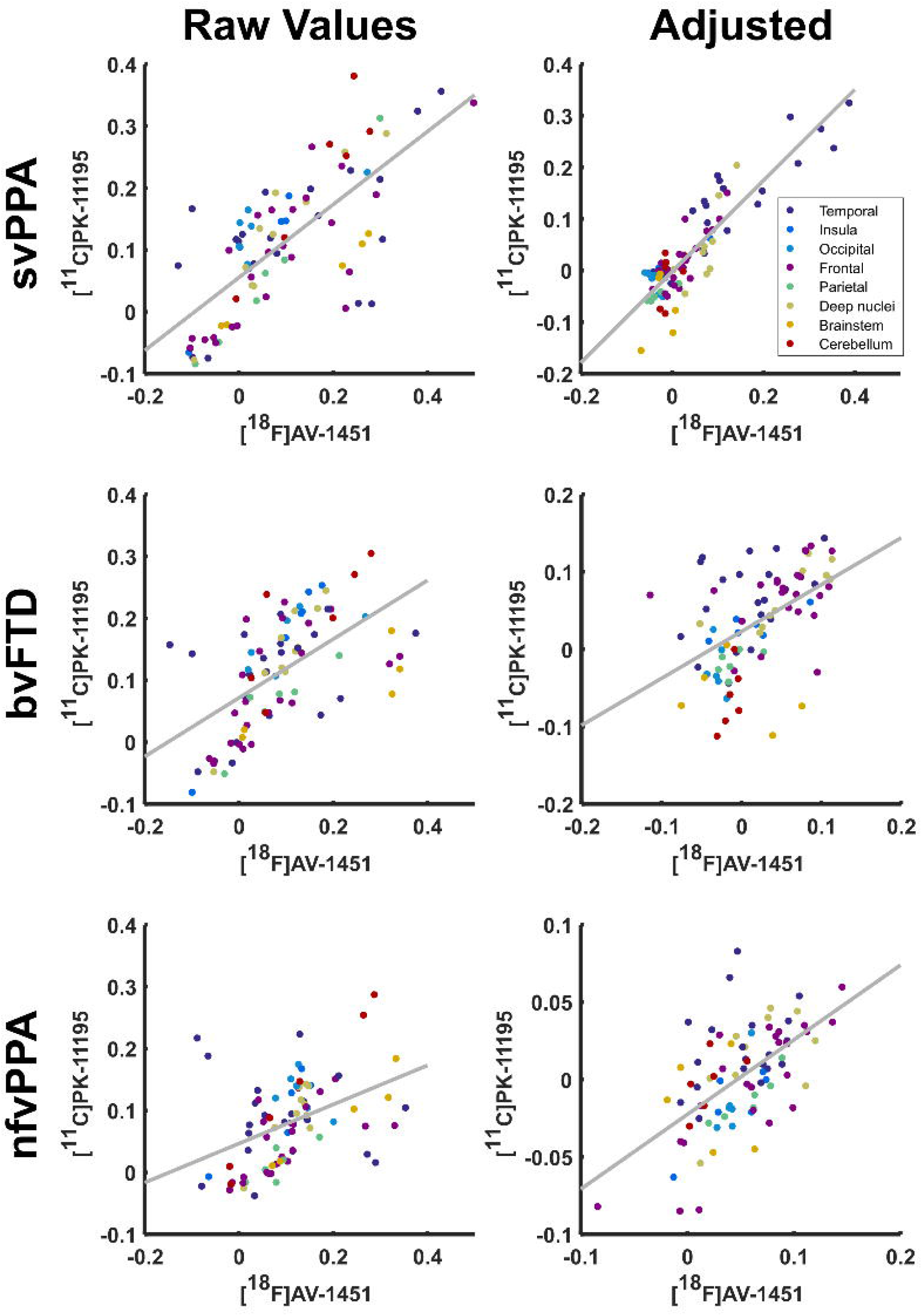
Scatter plot of the regional mean BP_ND_ for [^11^C]PK-11195 against regional mean BP_ND_ of [^18^F]AV-1451 by disease group. For each disease group raw values are demonstrated in the left hand plot with adjusted values demonstrated in the right hand plot.

### Principal component analysis of [^11^C]PK-11195 and [^18^F]AV-1451

Four principal components were detected in the [^11^C]-PK11195 BP_ND_ data before the elbow of the scree plot, which together explained 64% of the variance in the data (figure 3). Component 1 reflected whole brain binding. Component 2 was strongly weighted to the bilateral anterior temporal lobes. Component 3 primarily comprised frontal binding with a right sided predominance. Component 4 was not strongly loaded onto any single region but was weighted towards motor cortex. In a repeated measures ANOVA including these 3 principal components, there was a main effect of diagnosis (F(3, 39)=20.8, p<0.0001) and a significant interaction between principal component weighting and diagnosis (F(5.307, 68.99)=9.885, p<0.0001). *Post hoc* t-tests between individual disease groups and controls showed svPPA was associated with an increase in component 2 (t(10.2)=8.3, p<0.0001), bvFTD associated with both increased component 2 (t(9.297)=3.37, P=0.008) and component 3 (t(8.85)=3.95, p=0.003) and nfvPPA associated with increased component 3 (t(23)=2.68, p=0.013) (figure 3). Components 1 and 4 did not significantly differ between controls and patient groups.

**Figure 3:**
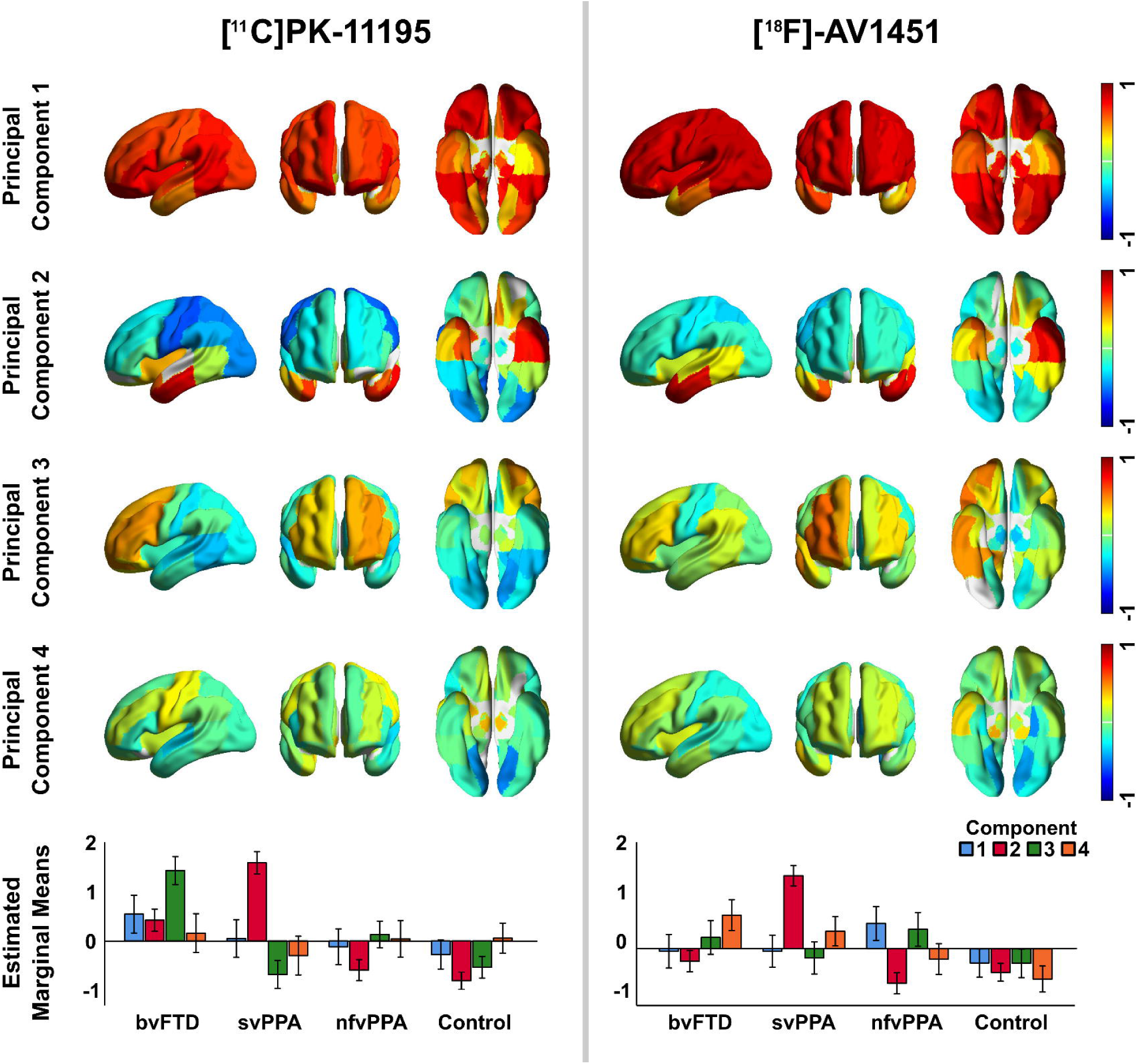
First four principal components for [^11^C]PK-11195 in the left column and [^18^F]AV-1451 in the right column. [^18^F]AV-1451 component 5 was also retained by Cattell’s criterion but was not strongly weighted to any region and did not discriminate groups so is omitted here for parsimony. The bottom row plots the estimated marginal means from the repeated measures ANOVA for each ligand, illustrating the association between principal component and diagnosis for each ligand.

Five principal components were detected in the [^18^F]AV-1451 BP_ND_ data before the elbow of the scree plot, which together accounted for 76% of the variance in the data. Component 1 again reflected global binding but less marked in the temporal poles, which were loaded onto component 2 (left) and component 4 (right). Component 3 was weighted asymmetrically towards frontal lobe binding. Component 5 was not strongly loaded onto any single region but was weighted towards bilateral superior temporal poles. In a repeated measures ANOVA including these 5 principal components, there was a main effect of diagnosis (F(3, 41)=5.43, p=0.003) and a significant interaction between principal component weighting and diagnosis (F(11, 150.6)=3.68, p<0.0001). The bvFTD group had increased weightings in component 3 (t(22)=2.345, p=0.28) and component 4 (t(11.575)=3.284, p=0.007), and svPPA had increased weightings in component 2 (t(15.005)=6.819, p<0.0001) and component 4 (t(12.9)=2.475, p=0.028) (figure 3). There were no significant *post hoc* differences between nfvPPA and controls. Components 1 and 5 did not significantly differ between controls and patient groups.

### Non-parametric analysis of [^11^C]PK-11195 and[^18^F]AV-1451 distributions

The principal component analyses suggest that a large amount of the variance between-subjects relates to whole brain PET signal. While this might reflect global differences in protein aggregation and neuroinflammation, it could also be explained by variations in radioligand affinity for different protein pathologies or other non-specific influences discussed below. We therefore performed an analysis of the relative *distribution* of PET signal for each individual scan, blinded to differences in overall signal magnitude by non-parametric rank-order statistical methods.

Multi-dimensional scaling plots of the non-parametric similarity between ligand distributions, for each sub-group pair and for all groups combined, are shown in figure 4. The CV-SVM classification accuracy and permutation-based statistical significance are indicated next to each plot. Classification was significantly better than chance in all cases, except for the finding that the non-parametric distribution of [^18^F]AV-1451 was unable to distinguish between bvFTD and nfvPPA.

**Figure 4:**
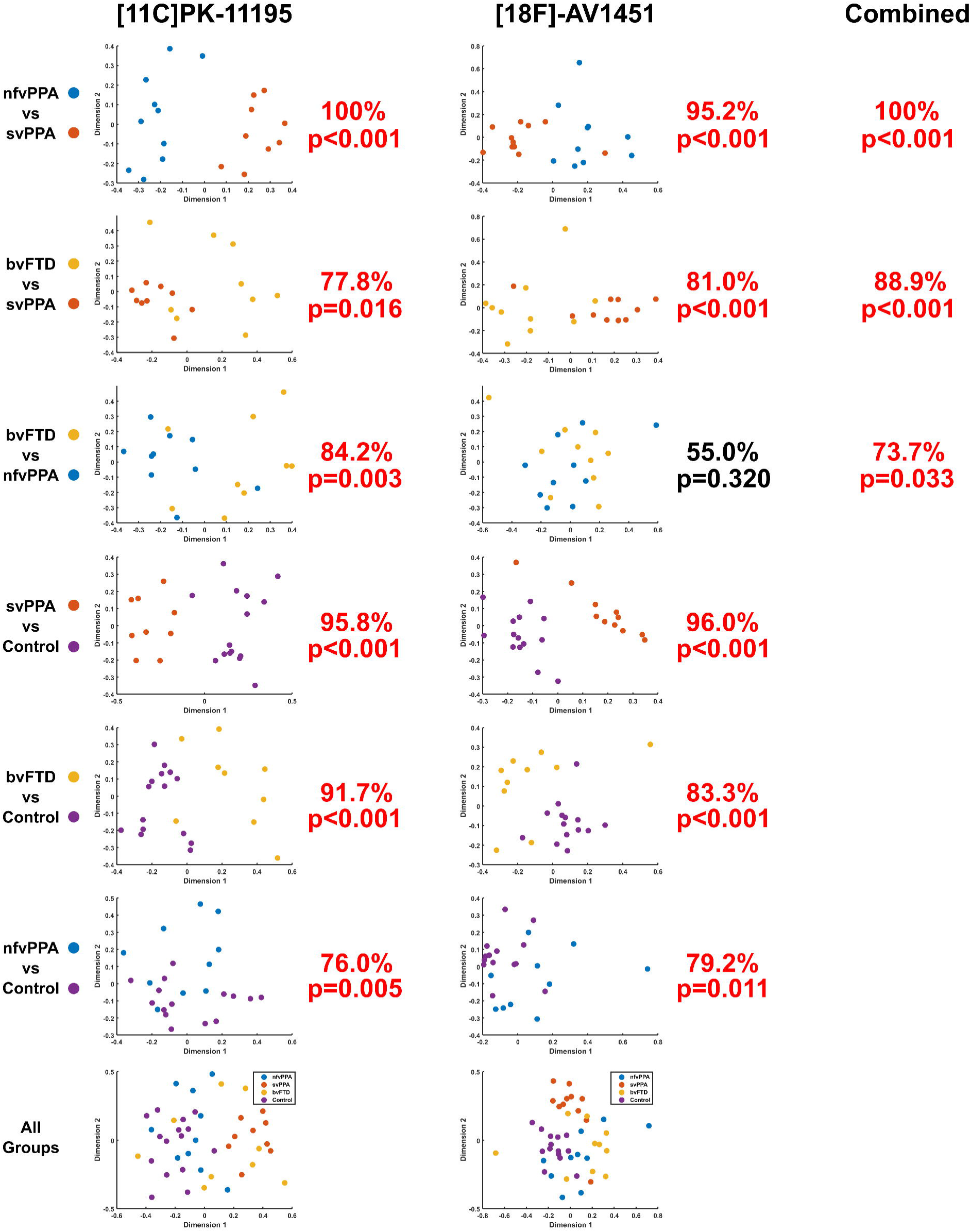
Pairwise classification accuracy for each ligand: [^11^C]PK-11195 on the left, [^18^F]AV-1451 in the middle, and using combined data on the right. The graphs represent a two-dimensional projection of the between-individual PET signal distribution dissimilarity according to the squared metric stress criterion. A ten-fold cross-validated support vector machine was applied to each plot, and the classification accuracy compared to a null distribution of 1000 randomisations for non-parametric significant testing. For each comparison percentage classification and p-value is stated.

For those FTD participants that underwent scanning with both ligands, the classification procedure was repeated after combining the multi-dimensional scaling data such that the CV-SVM operated on four dimensions rather than two. This resulted in an improvement in the differentiation of bvFTD and svPPA compared to either ligand alone (88.9% classification accuracy, p<0.001). Multimodal nfvPPA vs svPPA classification accuracy matched the performance of [^11^C]-PK11195 at 100%, p<0.001, but bvFTD vs nfvPPA classification performance was intermediate compared to each ligand alone, at 73.7%, p=0.033.

## Discussion

This *in vivo* study provides insights into complementary pathophysiological processes of frontotemporal dementia. Taken as a whole, our findings support an important role for neuroinflammation across the FTD spectrum, corroborating suggestions from epidemiological (Lant et al., 2014; Venneti et al., 2008), genetic (Broce et al., 2018; Guerreiro et al., 2013; Rayaprolu et al., 2013), imaging (Cagnin, Rossor, Sampson, MacKinnon, & Banati, 2004; Miyoshi et al., 2010) and animal studies (Bhaskar et al., 2010; Yin et al., 2010; Yoshiyama et al., 2007). Using regional analysis of variance, we have shown that neuroinflammation (indexed by [^11^C]PK-11195) and protein aggregates (Tau or TDP43, as indexed by [^18^F]AV-1451) are elevated across the FTD spectrum (figure 1). Furthermore, neuroinflammation is highly co-localised with protein aggregation within the individual syndromes, including most strongly in svPPA, where the predominant aggregated protein inclusions are TDP-43 rather than tau (figure 2). Principal component analysis also revealed distinct spatial modes of neuroinflammation, with frontotemporal, temporal pole and global distributions (figure 3). The weighting of these regional distributions differs between groups, supporting the regional differences in the pair-wise comparisons. The spatial modes of protein aggregation (figure 3) similarly reflect the well characterised distributions of pathology and are likewise weighted towards specific groups. However, the distribution of protein aggregation appears to be less focal than neuroinflammation in nfvPPA. To test the distinctiveness of inflammation and aggregation patterns, and to explicitly control for nonspecific, between-individual differences in ligand binding affinity, we used non-parametric multi-dimensional scaling and cross-validated linear support vector machines to classify patients. We demonstrated that the distribution of neuroinflammation can accurately distinguish each of the FTD syndromes from controls and from each other (figure 4). Classification was often possible based on the distribution of protein aggregation, but with less accuracy. The greater discriminatory ability of neuroinflammation emphasises its potential mechanistic relevance to the pathophysiology of FTD. Despite being strongly correlated at a regional level, the two PET tracers carry some unique information across these conditions, as illustrated by the improvement in distinguishing bvFTD from svPPA when multi-modal data were available to the classifier.

The correlation between regional distributions of neuroinflammation and protein aggregation supports a close relationship between these processes in FTD, mirroring recent evidence from Alzheimer’s disease that neuroinflammation is correlated with tau aggregation (Dani et al., 2018), and extending this to TDP-43 associated diseases. One interpretation of co-localised neuroinflammation and protein aggregation is that microglial activation is an early or initiating pathophysiological process, which promotes or accelerates abnormal protein misfolding and aggregation. Whilst it was previously thought that inflammation in the brain only occurred in the context of direct infection or after breakdown of the blood brain barrier, it is now recognised that microglia play a key role in orchestrating the innate immune response of the brain. They can be activated by misfolded proteins, and mediate responses through inflammatory pathways, cytotoxicity and changes in plasticity (Nakajima & Kohsaka, 2001; Nayak, Roth, & McGavern, 2014). In neurodegenerative diseases, this state of activation may become chronic, dysfunctional, and toxic, contributing to pathogenicity (Pasqualetti, Brooks, & Edison, 2015; Serrano-Pozo, Betensky, Frosch, & Hyman, 2016).

There is evidence for inflammatory processes in FTD (Heneka, Kummer, & Latz, 2014), from genetic (Broce et al., 2018; Guerreiro et al., 2013; Rayaprolu et al., 2013), cerebrospinal fluid (Sjogren et al., 2004; Woollacott et al., 2018), epidemiology (Miller et al., 2013, 2016), *post mortem* (Lant et al., 2014; Venneti et al., 2008) and animal studies (Bhaskar et al., 2010; Yin et al., 2010; Yoshiyama et al., 2007). It is well established that an innate immune response, characterised by activated microglia, is a feature of the neuropathology of FTD (Lant et al., 2014). Furthermore, mutations leading to haplo-insufficiency of progranulin, a growth factor that has peripheral immune and central microglial regulatory functions (Petkau et al., 2010; Pickford et al., 2011; Yin et al., 2010), leads to FTD syndromes characterised by TDP-43 pathology. Expansions in C9orf72 have effects on microglial function as well as neurons (O’Rouke et al., 2016), and risk variants for FTD in TREM2 are associated with microglial activation (Giraldo et al., 2013). Neuroinflammation is an early feature of pathophysiology in mouse models of tauopathy, where inflammatory changes precede the accumulation of aggregated tau (Yoshiyama et al., 2007) and pro-inflammatory molecules increase tau hyperphosphorylation and aggregation (Bhaskar et al., 2010). *In vivo* PET studies in small samples have shown that neuroinflammation anticipates atrophy in clinically established FTD (Cagnin et al., 2004) and precedes both symptoms and the detectability of tau aggregation by PET in MAPT mutation carriers (W Richard Bevan-Jones et al., 2018; Miyoshi et al., 2010). Although neuroinflammation appears early in the pathogenesis of FTD and other neurodegenerative disorders, it remains unclear whether it is an independently initiating factor or whether it is induced by oligomeric proteins or pre-tangles.

Much of the evidence supporting the presence of inflammation in FTD comes from *ex vivo* studies. The need to improve our understanding of this process during life has led to the development of PET radioligands for this purpose, but there is some controversy over the optimum ligand for imaging activated microglia. PET ligands which target TSPO have long been the mainstay of imaging microglia. However TSPO expression patterns in microglia are complex and the functional effects, i.e. deleterious versus protective, of different microglial phenotypes are incompletely understood (Gomez-Nicola & Perry, 2015). Furthermore, TSPO is also expressed by other cell types, notably astrocytes (McCarthy & Harden, 1981). However, in favour of the use of [^11^C]PK-11195 is its demonstrated selectivity for activated microglia over quiescent microglia and reactive astrocytes (Banati, 2002); its relative insensitivity to common polymorphisms in TSPO compared to second generation TSPO radioligands (Stefaniak & O’Brien, 2015; J. Zhang, 2015), and the fact it has well established methods of non-invasive kinetic analysis (L Passamonti et al., n.d.; Turkheimer et al., 2007). [^11^C]PK-11195 has also been effectively used in studies of other neurodegenerative diseases and shown ability to reveal pathologically-related patterns of neuroinflammation (Edison et al., 2013; L Passamonti et al., n.d.; Stefaniak & Brien, 2015; Varley, Brooks, & Edison, 2015). There remain some disadvantages, including relatively high non-specific binding and low brain penetration. Whilst this signal to noise has been cited as an explanation for previous negative studies using [^11^C]PK-11195, it does not undermine positive findings such as those shown here, especially within our multi-variate analyses that explicitly control for differences in ligand penetration and affinity. A further problem lies with interpreting the meaning of increased [^11^C]PK-11195 binding. This must include consideration of the potential contribution of reactive astrocytes expressing upregulated TSPO, but also our incomplete understanding of the functional consequences of activated microglia and reactive astrocytosis, and the potential effects of glial neuropathology on the immune/inflammatory component of pathophysiology. Whilst it seems reasonable to determine that increased [^11^C]PK-11195 binding equates to immune activation or neuroinflammation, the functional consequences or causality of this cannot be assumed.

In contrast to [^11^C]PK-11195, the [^18^F]AV-1451 binding provided a less clear signal despite such aggregation being an essential feature of FTD and many other dementias. We propose that [^18^F]AV-1451 binding is a proxy measure of aggregated non-β-amyloid protein. In Alzheimer’s disease the sensitivity of *in vivo* imaging with [^18^F]AV-1451, and its affinity for Tau in neurofibrillary tangles, is well established and has contributed significantly to our understanding of its pathogenesis and progression. However, the situation in FTD is more complex due to its pathological heterogeneity and our incomplete understanding of [^18^F]AV-1451 binding to the various morphologies of aggregated protein observed in FTLD-tau and FTLD-TDP43 pathologies. This heterogeneity is problematic and although six of our patients have genetic mutations, and six others were amyloid biomarker negative (tables 1&2), we cannot definitively ascertain the majority of patients’ pathological type *ante mortem.* In this context, the molecular targets for [^18^F]AV-1451 binding remain controversial. Supporting our use of [^18^F]AV-1451 as a marker of non-β-amyloid protein aggregation, previous *post mortem* work has demonstrated some binding to FTLD pathologies, albeit at a lower magnitude than that seen with Alzheimer’s pathology (Lowe et al., 2016; Marquié et al., 2015; Mcmillan et al., 2016; Sander et al., 2016). This appeared to be corroborated by *in vivo* studies of patients with a straight filament 4-repeat tauopathy and clinical FTD resulting from MAPT mutations, showing binding in areas typically affected in FTD and affected at *post mortem* (W. Richard Bevan-Jones et al., 2016; Smith et al., 2016), and by the elevated binding in the affected brain regions of patients with svPPA (William Richard Bevan-Jones et al., 2017; Makaretz et al., 2017) and bvFTD due to C9orf72 expansions (R. W. Bevan-Jones et al., 2018), who have TDP-43 rather than Tau pathology. However, even within genetically determined FTD binding affinity varies according to different Tau isoforms and strains (Jones et al., 2018) supporting varying affinity to different morphologies of Tau.

Other non-Tau, non-TDP-43, targets may also account for this increased binding. It is noted that this elevated [^18^F]AV-1451 binding is seen in a distribution that closely resembles that of neuropathology, suggesting potential binding to other proteins expressed by degenerating neurons or reactive glial cells, such as isoforms of monoamine oxidase (Vermeiren et al., 2018). Monoamine oxidase subtypes are expressed by both neurons (monoamine oxidase A) and reactive astrocytes (monoamine oxidase B) (Ben Haim, Carrillo-de Sauvage, Ceyzériat, & Escartin, 2015; Fowler, Logan, Volkow, & Wang, 2005) which may contribute to the patterns of cortical binding seen. Indeed, if [^18^F]AV-1451 binding were driven by ‘off target’ binding to reactive astrocytes, which are induced by activated microglia (Liddelow et al., 2017), this would only provide further evidence for the importance of neuroinflammation in FTD. Further to this, upregulation of TSPO in reactive astrocytes could occur alongside that of MAO-B accounting for the regional correlation found between ligand binding. However, the pre-symptomatic dissociation of [^11^C]PK-11195 and [^18^F]AV-1451 binding (W Richard Bevan-Jones et al., 2018) argues strongly against such simple cross-affinity.

In the face of uncertainty about molecular targets and variations in affinity, it is important to emphasise that through our classification analysis we focus on distribution rather than quantification of binding, using a non-parametric method that is insensitive to absolute binding values and purely reflects the pattern of binding. This takes into account the potential differences in affinity of [^18^F]AV-1451 for different protein targets. Overall, whilst it is clear that [^18^F]AV-1451 binding does not bind specifically to Tau aggregates, the distribution of binding co-localises and varies with that expected of aggregated protein in these diseases, and *post mortem* immunohistochemistry of Tau. Indeed, [^18^F]AV-1451 may provide a usefully non-selective marker of non-βamyloid aggregated protein, whether Tau or TDP-43, allowing *in vivo* examination across the spectrum of sporadic FTD syndromes. Whilst in the complex setting of FTLD we interpret [^18^F]AV-1451 binding as a non-specific marker of non-β-amyloid neuropathology, the biological relevance of elevated binding in non-AD neurodegenerative disease remains incompletely understood. Further work examining [^18^F]AV-1451 binding across large post mortem cohorts of FTLD pathology will be required to independently validate our hypothesis.

The main limitation of this study is group size which, although larger than most previous PET studies in FTD, is still small for each individual diagnosis. The small sample size reduces the power of the study to find parametric group differences in binding, particularly given that both ligands have a degree of insensitivity to their target, as well as limiting the ability to detect associations with clinical features and severity. Characterisation of the groups is also limited in that the genotyping and amyloid assays were based on clinical indications and consent: we did not directly examine amyloid status in all individuals and whilst there is a mix of both genetic and sporadic cases we did not genotype every participant. The inability to perform pathological subtyping *in vivo* makes interpretation of results more difficult in view of the generally poor relationship between phenotype and underlying neuropathology in FTD. Consequently, we cannot use the clinical diagnostic groups alone to draw conclusions about the relationship between microglial activation and specific forms of protein aggregation. We also are limited in the inferences about the predilection for immune dysregulation in a particular neuropathological subtype, such as the relationship suggested between immune dysfunction and FTLD-TDP-43 (Miller et al., 2013, 2016), except for the cases with genetic mutations.

To conclude, we provide *in vivo* evidence for neuroinflammation in FTD, which has a close relationship with [^18^F]AV-1451 binding, taken in this study to represent a marker of FTLD-tau or FTLD-TDP-43 neuropathology. PET measurement of inflammation provided a more accurate classification of syndromes than did protein aggregation emphasising its potential importance in shaping the clinical and neuropathological patterns of the diverse clinical syndromes of frontotemporal dementia. A causal role for neuroinflammation in neurodegeneration would inform future drug targets and potential clinical trials in frontotemporal dementia. Our findings therefore warrant further longitudinal mechanistic investigation into the role of neuroinflammation in early-stage neurodegeneration, its relationship to specific protein aggregation and to clinical progression.

## Supporting information

weak correlations were observed between our cohorts of controls for each ligand (supplementary figure 1)

## Acknowledgements

We thank our volunteers for participating in this study and to the radiographers/technologists at the Wolfson Brain Imaging Centre and PET/CT Unit, Addenbrooke’s Hospital, for their invaluable support in data acquisition. We thank the NIHR Dementias and Neurodegenerative Diseases Research Network for help with subject recruitment. We also thank Dr’s Istvan Boros, Joong-Hyun Chun, and WBIC RPU for the manufacture of the radioligands. We thank Avid (Lilly) for supplying the precursor for the manufacturing of [^18^F]AV-1451 for use in this study. The work was supported by National Institute for Health Research Cambridge Biomedical Research Centre; the Wellcome Trust (JBR 103838); the Association of British Neurologists and the Patrick Berthoud Charitable Trust (TEC).

---

***Supplementary figure 1:*** Scatter plot of the raw regional mean BP_ND_ for [^11^C]PK-11195 against regional mean BP_ND_ of [^18^F]AV-1451 between the control groups.

## References

Banati, R. B. (2002). Visualising Microglial Activation In Vivo. Glia, (February), 206–217. http://doi.org/10.1002/glia.10144

Ben Haim, L., Carrillo-de Sauvage, M.-A., Ceyzériat, K., & Escartin, C. (2015). Elusive roles for reactive astrocytes in neurodegenerative diseases. Frontiers in Cellular Neuroscience, 9(August), 1–27. http://doi.org/10.3389/fncel.2015.00278

Bevan-Jones, R. W., Cope, T. E., Jones, S. P., Passamonti, L., Hong, Y. T., Fryer, T., … Rowe, J. B. (2018). [18F]AV-1451 binding is increased in frontotemporal dementia due to C9orf72 expansion. Annals of Clinical and Translational Neurology, 11–13. http://doi.org/10.1002/acn3.631

Bevan-Jones, W. R., Cope, T. E., Jones, P. S., Passamonti, L., Hong, Y. T., Fryer, T. D., … Rowe, J. B. (2017). [18F] AV-1451 binding in vivo mirrors the expected distribution of TDP-43 pathology in the semantic variant of primary progressive aphasia. Journal of Neurology Neurosurgery and Psychiatry, 1–6. http://doi.org/10.1136/jnnp-2017-316402

Bevan-Jones, W. R., Cope, T. E., Jones, P. S., Passamonti, L., Hong, Y. T., Fryer, T., … Rowe, J. B. (2018). In vivo evidence for pre-symptomatic neuroinflammation in a MAPT mutation carrier. Annals of Clinical and Translational Neurology, in press.

Bevan-Jones, W. R., Cope, T. E., Passamonti, L., Fryer, T. D., Hong, Y. T., Aigbirhio, F., … Rowe, J. B. (2016). [18F]AV-1451 PET in behavioral variant frontotemporal dementia due to MAPT mutation. Annals of Clinical and Translational Neurology, 3(12), 940–947. http://doi.org/10.1002/acn3.366

Bevan-Jones, W. R., Surendranathan, A., Passamonti, L., Rodríguez, P. V., Arnold, R., Mak, E., … Brien, J. T. O. (2017). Neuroimaging of Inflammation in Memory and Related Other Disorders (NIMROD) study protocol: a deep phenotyping cohort study of the role of brain inflammation in dementia, depression and other neurological illnesses. BMJ Open, 7(1), 1–9. http://doi.org/10.1136/bmjopen-2016-013187

Bhaskar, K., Konerth, M., Kokiko-cochran, O. N., Cardona, A., Ransohoff, R. M., & Lamb, B. T. (2010). Regulation of Tau Pathology by the Microglial Fractalkine Receptor. Neuron, 68(1), 19–31. http://doi.org/10.1016/j.neuron.2010.08.023|

Broce, I., Karch, C. M., Wen, N., Fan, C. C., Wang, Y., Hong, C., … Sugrue, L. P. (2018). Immune-related genetic enrichment in frontotemporal dementia[]: An analysis of genome-wide association studies. PLOS Medicine, 1–20.

Cagnin, A., Rossor, M., Sampson, E. L., MacKinnon, T., & Banati, R. B. (2004). In Vivo Detection of Microglial Activation in Frontotemporal Dementia. Annals of Neurology, 56(6), 894–897. http://doi.org/10.1002/ana.20332

Chien, D. T., Bahri, S., Szardenings, A. K., Walsh, J. C., & Mu, F. (2013). Early Clinical PET Imaging Results with the Novel PHF-Tau Radioligand [F-18]-T807. Journal of Alzheimer’s Disease, 34, 457–468. http://doi.org/10.3233/JAD-122059

Dani, M., Wood, M., Mizoguchi, R., Fan, Z., Walker, Z., Morgan, R., … Edison, P. (2018). Microglial activation correlates in vivo with both tau and amyloid in Alzheimer’s disease. Brain, (July), 1–15. http://doi.org/10.1093/brain/awy188

Edison, P., Ahmed, I., Fan, Z., Hinz, R., Gelosa, G., Ray Chaudhuri, K., … Brooks, D. J. (2013). Microglia, amyloid, and glucose metabolism in Parkinson’s disease with and without dementia. Neuropsychopharmacology, 38(6), 938–49. http://doi.org/10.1038/npp.2012.255

Fowler, J. S., Logan, J., Volkow, N. D., & Wang, G. (2005). Translational Neuroimaging[]: Positron Emission Tomography Studies of Monoamine Oxidase. Molecular Imaging and Biology, 7(November), 377–387. http://doi.org/10.1007/s11307-005-0016-1

Giraldo, M., Lopera, F., Siniard, A. L., Corneveaux, J. J., Schrauwen, I., Carvajal, J., … Huentelman, M. J. (2013). Variants in triggering receptor expressed on myeloid cells 2 are associated with both behavioral variant frontotemporal lobar degeneration and Alzheimer’s disease. Neurobiology of Aging, 34(8), 2077.e11-2077.e18. http://doi.org/10.1016/j.neurobiolaging.2013.02.016

Gomez-Nicola, D., & Perry, V. H. (2015). Microglial Dynamics and Role in the Healthy and Diseased Brain[]: A Paradigm of Functional Plasticity. The Neuroscientist, 21(2), 169–184. http://doi.org/10.1177/1073858414530512

Gorno-Tempini, M. L., Hillis, A. E., Weintraub, S., Kertesz, A., Mendez, M., Cappa, S. F., … Grossman, M. (2011). Classification of primary progressive aphasia and its variants. Neurology, 76(11), 1006–14. http://doi.org/10.1212/WNL.0b013e31821103e6

Guerreiro, R. J., Lohmann, E., Bras, J. M., Gibbs, J. R., Rohrer, J. D., Gurunlian, N., … Hardy, J. (2013). Using Exome Sequencing to Reveal Mutations in TREM2 Presenting as a Frontotemporal Dementia-like Syndrome Without Bone Involvement. JAMA Neurology, 70(1), 78–84. http://doi.org/10.1001/jamaneurol.2013.579

Heneka, M. T., Kummer, M. P., & Latz, E. (2014). Innate immune activation in neurodegenerative disease. Nature Reviews Immunology, 14(July), 463–477. http://doi.org/10.1038/nri3705

Jones, D. T., Knopman, D. S., Graff-Radford, J., Syrjanen, J. A., Senjem, M. L., Schwarz, C. G., … Boeve, B. F. (2018). In vivo F-AV-1451 tau-PET signal in MAPT mutation carriers varies by expected tau isoforms. Neurology, (February). http://doi.org/10.1212/WNL.0000000000005117

Kriegeskorte, N., Mur, M., & Bandettini, P. (2008). Representational similarity analysis – connecting the branches of systems neuroscience. Frontiers in Systems Neuroscience, 2(4), 1–28. http://doi.org/10.3389/neuro.06.004.2008

Lant, S. B., Robinson, A. C., Thompson, J. C., Rollinson, S., Pickering-Brown, S., Snowden, J. S., … Neurobiology, A. (2014). Patterns of microglial cell activation in frontotemporal lobar degeneration. Neuropathology and Applied Neurobiology, 40, 686–696. http://doi.org/10.1111/nan.12092

Liddelow, S. A., Guttenplan, K. A., Clarke, L. E., Bennett, F. C., Bohlen, C. J., Schirmer, L., … Barres, B. A. (2017). Neurotoxic reactive astrocytes are induced by activated microglia. Nature, 541(7638), 481–487. http://doi.org/10.1038/nature21029

Lowe, V. J., Curran, G., Fang, P., Liesinger, A. M., Josephs, K. A., Parisi, J. E., … Murray, M. E. (2016). An autoradiographic evaluation of AV-1451 Tau PET in dementia. Acta Neuropathologica Communications, 4(1), 58. http://doi.org/10.1186/s40478-016-0315-6

MacKenzie, I. R. A., Neumann, M., Bigio, E. H., Cairns, N. J., Alafuzoff, I., Kril, J., … Mann, D. M. A. (2010). Nomenclature and nosology for neuropathologic subtypes of frontotemporal lobar degeneration: An update. Acta Neuropathologica, 119(1), 1–4. http://doi.org/10.1007/s00401-009-0612-2

Makaretz, S. J., Quimby, M., Collins, J., Makris, N., Mcginnis, S., Schultz, A., … Dickerson, B. C. (2017). Flortaucipir tau PET imaging in semantic variant primary progressive aphasia. Journal of Neurology Neurosurgery and Psychiatry, 1–8. http://doi.org/10.1136/jnnp-2017-316409

Marquié, M., Normandin, M. D., Vanderburg, C. R., Costantino, I. M., Bien, E. A., Rycyna, L. G., … Gómez-Isla, T. (2015). Validating novel tau positron emission tomography tracer [F-18]-AV-1451 (T807) on postmortem brain tissue. Annals of Neurology, 78(5), 787–800. http://doi.org/10.1002/ana.24517

McCarthy, K. D., & Harden, T. K. (1981). Identification of two benzodiazepine binding sites on cells cultured from rat cerebral cortex. J.Pharmacol.Exp.Ther., 216, 183–191.

Mcmillan, C. T., Irwin, D. J., Nasrallah, I., Phillips, J. S., Spindler, M., Rascovsky, K., … Grossman, M. (2016). Multimodal evaluation demonstrates in vivo 18F-AV-1451 uptake in autopsy-confirmed corticobasal degeneration. Acta Neuropathologica, 132(6), 935–937. http://doi.org/10.1007/s00401-016-1640-3

Miller, Z. A., Rankin, K. P., Graff-radford, N. R., Takada, L. T., Sturm, V. E., Cleveland, C. M., … Miller, B. L. (2013). TDP-43 frontotemporal lobar degeneration and autoimmune disease. Jornal of Neurology, Neurosurgery and Psychiatry, 84, 956–962. http://doi.org/10.1136/jnnp-2012-304644

Miller, Z. A., Sturm, V. E., Camsari, G. B., Karydas, A., Yokoyama, J. S., Grinberg, L. T., … Miller, B. L. (2016). Increased prevalence of autoimmune disease within C9 and FTD/MND cohorts Completing the picture. Neurology: Neuroimmunology and NeuroInflammation, 3(6), 1–9. http://doi.org/10.1212/NXI.0000000000000301

Miyoshi, M., Shinotoh, H., Wszolek, Z. K., Strongosky, A. J., Shimada, H., Arakawa, R., … Suhara, T. (2010). In vivo detection of neuropathologic changes in presymptomatic MAPT mutation carriers[]: A PET and MRI study. Parkinsonism and Related Disorders, 16(6), 404–408. http://doi.org/10.1016/j.parkreldis.2010.04.004

Nakajima, K., & Kohsaka, S. (2001). Microglia: Activation and Their Significance in the Central Nervous System. Journal of Biochemistry, 130(2), 169–175.

Nayak, D., Roth, T. L., & McGavern, D. B. (2014). Microglia development and function. Annual Review of Immunology, 32, 367–402. http://doi.org/10.1146/annurev-immunol-032713-120240

O’Rouke, J. G., Bogdanik, L., Yanez, A., Lall, D., Wolf, A. J., Muhammad, A. K. M. G., … Baloh, R. H. (2016). C9orf72 is required for proper macrophage and microglial function in mice. Science, 351(6279).

Pasqualetti, G., Brooks, D. J., & Edison, P. (2015). The Role of Neuroinflammation in Dementias. Current Neurology and Neuroscience Reports, 15. http://doi.org/10.1007/s11910-015-0531-7

Passamonti, L., Vazquez Rodriguez, P., Hong, Y. T., Allinson, K. S. J., Bevan-Jones, W. R., Williamson, D., … Rowe, J. B. (n.d.). [11C]PK11195 binding in Alzheimer’s disease and progressive supranuclear palsy. Neurology, 1–20.

Passamonti, L., Vazquez Rodriguez, P., Hong, Y. T., Allinson, K. S. J., Williamson, D., Borchert, R. J., … Rowe, J. B. (2017). 18F-AV-1451 positron emission tomography in Alzheimer’s disease and progressive supranuclear palsy. BrainfZ: A Journal of Neurology, 140(January), 1–11. http://doi.org/10.1093/brain/aww340

Petkau, T. L., Neal, S. J., Orban, P. C., Macdonald, J. L., Hill, A. M., Lu, G., … Leavitt, B. R. (2010). Progranulin Expression in the Developing and Adult Murine Brain. Research in Systems Neuroscience, 518, 3931–3947. http://doi.org/10.1002/cne.22430

Pickford, F., Marcus, J., Camargo, L. M., Xiao, Q., Graham, D., Mo, J., … Hutton, M. (2011). Progranulin Is a Chemoattractant for Microglia and Stimulates Their Endocytic Activity. The American Journal of Pathology, 178(1), 284–295. http://doi.org/10.1016/j.ajpath.2010.11.002

Rascovsky, K., Hodges, J. R., Knopman, D., Mendez, M. F., Kramer, J. H., Neuhaus, J., … Miller, B. L. (2011). Sensitivity of revised diagnostic criteria for the behavioural variant of frontotemporal dementia. Brain, 134(9), 2456–2477. http://doi.org/10.1093/brain/awr179

Rayaprolu, S., Mullen, B., Baker, M., Lynch, T., Finger, E., Seeley, W. W., … Ross, O. A. (2013). TREM2 in neurodegeneration[]: evidence for association of the p. R47H variant with frontotemporal dementia and Parkinson’s disease. Molecular Neurodegeneration, 8, 1–5.

Rogalski, E., Cobia, D., Harrison, T. M., Wieneke, C., Thompson, C. K., Weintraub, S., & Mesulam, M.-M. (2011). Anatomy of Language Impairments in Primary Progressive Aphasia. Journal of Neuroscience, 31(9), 3344–3350. http://doi.org/10.1523/JNEUROSCI.5544-10.2011

Sander, K., Lashley, T., Gami, P., Gendron, T., Lythgoe, M. F., Rohrer, J. D., … ??rstad, E. (2016). Characterization of tau positron emission tomography tracer [18F]AV-1451 binding to postmortem tissue in Alzheimer’s disease, primary tauopathies, and other dementias. Alzheimer’s and Dementia, 12(11), 1116–1124. http://doi.org/10.1016/j.jalz.2016.01.003

Seelaar, H., Rohrer, J. D., Pijnenburg, Y. A. L., Fox, N. C., & van Swieten, J. C. (2011). Clinical, genetic and pathological heterogeneity of frontotemporal dementia: a review. Journal of Neurology, Neurosurgery & Psychiatry, 82(5), 476–486. http://doi.org/10.1136/jnnp.2010.212225

Serrano-Pozo, A., Betensky, R. A., Frosch, M. P., & Hyman, B. T. (2016). Plaque-Associated Local Toxicity Increases over the Clinical Course of Alzheimer Disease. The American Journal of Pathology, 186(2), 375–384. http://doi.org/10.1016/j.ajpath.2015.10.010

Sjogren, M., Folkesson, S., Blennow, K., & Tarkowski, E. (2004). Increased intrathecal inflammatory activity in frontotemporal dementia: pathophysiological implications. Journal of Neurology Neurosurgery and Psychiatry, 75, 1107–1111. http://doi.org/10.1136/jnnp.2003.019422

Smith, R., Puschmann, A., Scholl, M., Ohlsson, T., Van Swieten, J., Honer, M., … Hansson, O. (2016). 18F-AV-1451 tau PET imaging correlates strongly with tau neuropathology in MAPT mutation carriers. Brain, 139(9), 2372–2379. http://doi.org/10.1093/brain/aww163

Smith, R., Scholl, M., Widner, H., van Westen, D., Svenningsson, P., Hägerström, D., … Hansson, O. (2017). In vivo retention of 18 F-AV-1451 in corticobasal syndrome. Neurology, 1–10.

Spinelli, E. G., Mandelli, M. L., Miller, Z. A., Santos-, M. A., Wilson, S. M., Agosta, F., … Gorno-tempini, M. L. (2017). Typical and atypical pathology in primary progressive aphasia variants. Annals of Neurology, 81(3), 430–443.

Stefaniak, J., & Brien, J. O. (2015). Imaging of neuroinflammation in dementia: a review. Journal of Neurology Neurosurgery and Psychiatry, 1–8. http://doi.org/10.1136/jnnp-2015-311336

Stefaniak, J., & O’Brien, J. (2015). Imaging of neuroinflammation in dementia: a review. Journal of Neurology, Neurosurgery & Psychiatry, 21–28. http://doi.org/10.1136/jnnp-2015-311336

Turkheimer, F. E., Edison, P., Pavese, N., Roncaroli, F., Anderson, A. N., Hammers, A., … Brooks, D. J. (2007). Reference and Target Region Modeling of [11C]-(R)-PK11195 Brain Studies. The Journal of Nuclear Medicine, 48(1), 158–167.

Varley, J., Brooks, D. J., & Edison, P. (2015). Imaging neuroinflammation in Alzheimer’s disease and other dementias[]: Recent advances and future directions. Alzheimer’s & Dementia, 11(9), 1110–1120. http://doi.org/10.1016/j.jalz.2014.08.105

Venneti, S., Wang, G., Nguyen, J., & Wiley, C. A. (2008). The Positron Emission Tomography Ligand DAA1106 Binds With High Affinity to Activated Microglia in Human Neurological Disorders. Journal of Neuropathology and Experimental Neurology, 67(10), 1001–1010.

Vermeiren, C., Motte, P., Viot, D., Mairet-Coello, G., Courade, J.-P., Citron, M., … Gillard, M. (2018). The Tau Positron-Emission Tomography Tracer AV-1451 Binds With Similar Affinities to Tau Fibrils and Monoamine Oxidases. Movement Disorders, 33(2), 273–281. http://doi.org/10.1002/mds.27271

Woollacott, I. O. C., Nicholas, J. M., Heslegrave, A., Heller, C., Foiani, M. S., Dick, K. M., … Rohrer, J. D. (2018). Cerebrospinal fluid soluble TREM2 levels in frontotemporal dementia differ by genetic and pathological subgroup. Alzheimer’s Research and Therapy, 10(1), 1–14. http://doi.org/10.1186/s13195-018-0405-8

Xia, C., Arteaga, J., Chen, G., Gangadharmath, U., Gomez, L. F., Kasi, D., … Kolb, H. C. (2013). [18 F] T807, a novel tau positron emission tomography imaging agent for Alzheimer’s disease. Alzheimer’s & Dementia, 9(6), 666–676. http://doi.org/10.1016/j.jalz.2012.11.008

Yaqub, M., van Berckel, B. N. M., Schuitemaker, A., Hinz, R., Turkheimer, F. E., Tomasi, G., … Boellaard, R. (2012). Optimization of supervised cluster analysis for extracting reference tissue input curves in (R)-[(11)C]PK11195 brain PET studies. Journal of Cerebral Blood Flow and MetabolismfZ: Official Journal of the International Society of Cerebral Blood Flow and Metabolism, 32(8), 1600–8. http://doi.org/10.1038/jcbfm.2012.59

Yin, F., Banerjee, R., Thomas, B., Zhou, P., Qian, L., Jia, T., … Ding, A. (2010). Exaggerated inflammation, impaired host defense, and neuropathology in progranulin-deficient mice. The Journal of Experimental Medicine, 207(1), 117–128. http://doi.org/10.1084/jem.20091568

Yoshiyama, Y., Higuchi, M., Zhang, B., Huang, S., Iwata, N., Saido, T. C., … Lee, V. M. (2007). Synapse Loss and Microglial Activation Precede Tangles in a P301S Tauopathy Mouse Model. Neuron, 53, 337–351. http://doi.org/10.1016/j.neuron.2007.01.010

Zhang, J. (2015). Mapping neuroinflammation in frontotemporal dementia with molecular PET imaging. J Neuroinflammation, 12, 108. http://doi.org/10.1186/s12974-015-0236-5

Zhang, W., Arteaga, J., Cashion, D. K., Chen, G., Gangadharmath, U., Gomez, L. F., … Xia, C. (2012). A Highly Selective and Specific PET Tracer for Imaging of Tau Pathologies. Journal of Alzheimer’s Disease, 31(3), 601–612. http://doi.org/10.3233/JAD-2012-120712

