## Supplementary figures and images for "Neuroinflammation and protein aggregation co-localize across the frontotemporal dementia spectrum"

### weak correlations were observed between our cohorts of controls for each ligand (supplementary figure 1)

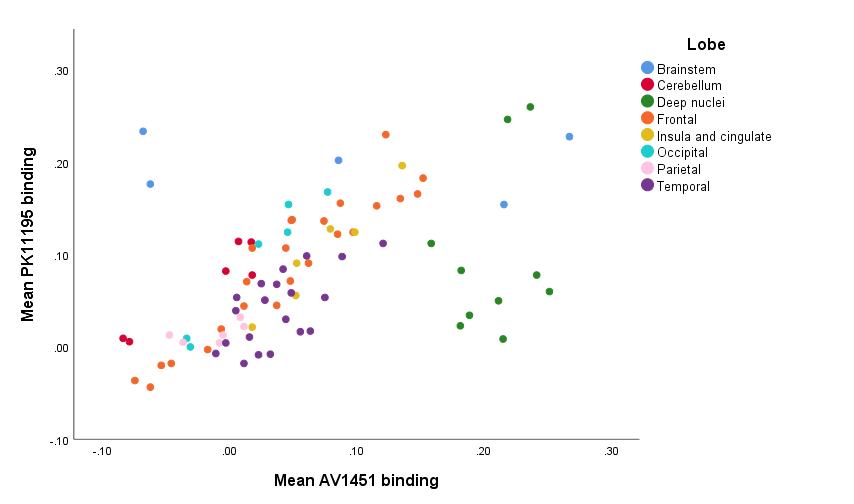
